# The *C. elegans* SMOC-1 protein acts cell non-autonomously to promote bone morphogenetic protein signaling

**DOI:** 10.1101/416669

**Authors:** Melisa S. DeGroot, Herong Shi, Alice Eastman, Alexandra N. McKillop, Jun Liu

## Abstract

Bone morphogenetic protein (BMP) signaling regulates many different developmental and homeostatic processes in metazoans. The BMP pathway is conserved in *Caenorhabditis elegans*, and is known to regulate body size and mesoderm development. We have identified the *C. elegans smoc-1* (Secreted MOdular Calcium binding protein-1) gene as a new player in the BMP pathway. *smoc-1(0)* null mutants have a small body size, while overexpression of *smoc-1* led to a long body size and increased expression of the RAD-SMAD BMP reporter, suggesting that SMOC-1 acts as a positive modulator of BMP signaling. Using double mutant analysis, we showed that SMOC-1 antagonizes the function of the glypican LON-2 and acts through the BMP ligand DBL-1 to regulate BMP signaling. Moreover, SMOC-1 appears to specifically regulate BMP signaling without significant involvement in a TGFβ-like pathway that regulates dauer development. We found that *smoc-1* is expressed in multiple tissues, including cells of the pharynx, intestine, and posterior hypodermis, and that the expression of *smoc-1* in the intestine is positively regulated by BMP signaling. We further established that SMOC-1 functions cell non-autonomously to regulate body size. Human SMOC1 and SMOC2 can each partially rescue the *smoc-1(0)* mutant phenotype, suggesting that SMOC-1’s function in modulating BMP signaling is evolutionarily conserved. Together, our findings highlight a conserved role of SMOC proteins in modulating BMP signaling in metazoans.

**ARTICLE SUMMARY:** BMP signaling is critical for development and homeostasis in metazoans, and is under tight regulation. We report the identification and characterization of a Secreted MOdular Calcium binding protein SMOC-1 as a positive modulator of BMP signaling in *C. elegans*. We established that SMOC-1 antagonizes the function of LON-2/glypican and acts through the DBL-1/BMP ligand to promote BMP signaling. We identified *smoc-1*-expressing cells, and demonstrated that SMOC-1 acts cell non-autonomously and in a positive feedback loop to regulate BMP signaling. We also provide evidence suggesting that the function of SMOC proteins in the BMP pathway is conserved from worms to humans.

## INTRODUCTION

Bone morphogenetic proteins (BMPs) are highly conserved signaling molecules that mediate cell-cell communication. The BMP signaling cascade is initiated when the BMP ligands bind to the membrane-bound receptor kinases, upon which the type-II receptor phosphorylates the type-I receptors. The signaling cascade is then transduced within the receiving cell as the receptor-associated Smads (R-Smads) are activated via phosphorylation by the type-I receptor. Activated R-Smads complex together with common mediator Smads (co-Smads) and other transcription factors to regulate transcription of downstream genes (KATAGIRI AND WATABE 2016). BMPs regulate fundamental cellular processes, including cell migration, cell proliferation, cell fate specification, and cell death throughout metazoan development (WANG *et al.* 2014). Tight regulation of BMP signaling in time, space, magnitude, and duration is therefore important for proper developmental outcomes. Mis-regulation of BMP signaling can cause a variety of disorders in humans (BRAZIL *et al.* 2015; SALAZAR *et al.* 2016; WU *et al.* 2016). Previous studies have demonstrated that BMP signaling can be regulated at many levels, both extracellularly and intracellularly (BRAGDON *et al.* 2011; LOWERY *et al.* 2016; SEDLMEIER AND SLEEMAN 2017). The nematode *C. elegans* provides a useful system for identifying factors that modulate the BMP pathway.

The BMP pathway in *C. elegans* is comprised of evolutionarily conserved core components including the ligand (DBL-1/BMP), the type I and type II receptors (SMA-6/RI and DAF-4/RII), the R-Smads (SMA-2 and SMA-3), and the co-Smad (SMA-4) (ESTEVEZ *et al.* 1993; SAVAGE *et al.* 1996; KRISHNA *et al.* 1999; MORITA *et al.* 1999; SUZUKI *et al.* 1999; MORITA *et al.* 2002) (Figure 1A). Unlike in *Drosophila* and vertebrates, BMP signaling is not essential for viability in *C. elegans*, yet it regulates multiple processes, including body size, male tail development, and mesoderm patterning (GUMIENNY AND SAVAGE-DUNN 2013; SAVAGE-DUNN AND PADGETT 2017). The BMP ligand DBL-1 is expressed in the ventral nerve cord (SUZUKI *et al.* 1999), and it activates the pathway in the hypodermis to regulate body size (YOSHIDA *et al.* 2001; WANG *et al.* 2002). Reduced BMP signaling causes a small (Sma) body size, while increased BMP signaling leads to a long (Lon) body size (MORITA *et al.* 1999; SUZUKI *et al.* 1999; MORITA *et al.* 2002). BMP signaling also regulates the development of the postembryonic mesoderm lineage, the M lineage. We have shown that mutations in the BMP pathway specifically suppress the M lineage dorsoventral patterning defects caused by mutations in *sma-9*, which encodes the *C. elegans* zinc finger protein Schnurri (LIANG *et al.* 2003; FOEHR *et al.* 2006). Specifically, mutations in *sma-9* result in the loss of the two M-derived coelomocytes (CCs), while BMP pathway mutations can restore these two CCs in the *sma-9(0)* mutant background (FOEHR *et al.* 2006; LIU *et al.* 2015; WANG *et al.* 2017) (Figure 1B,C). Using this suppression of *sma-9(0)* M-lineage defect (Susm) assay, we have identified multiple evolutionarily conserved modulators of BMP signaling. These include the RGM protein DRAG-1 (TIAN *et al.* 2010), the neogenin homolog UNC-40 (TIAN *et al.* 2013), the ADAM10 protein SUP-17 (WANG *et al.* 2017), and three tetraspanins, TSP-21, TSP-12 and TSP-14 (LIU *et al.* 2015; WANG *et al.* 2017).

**Fig 1.**
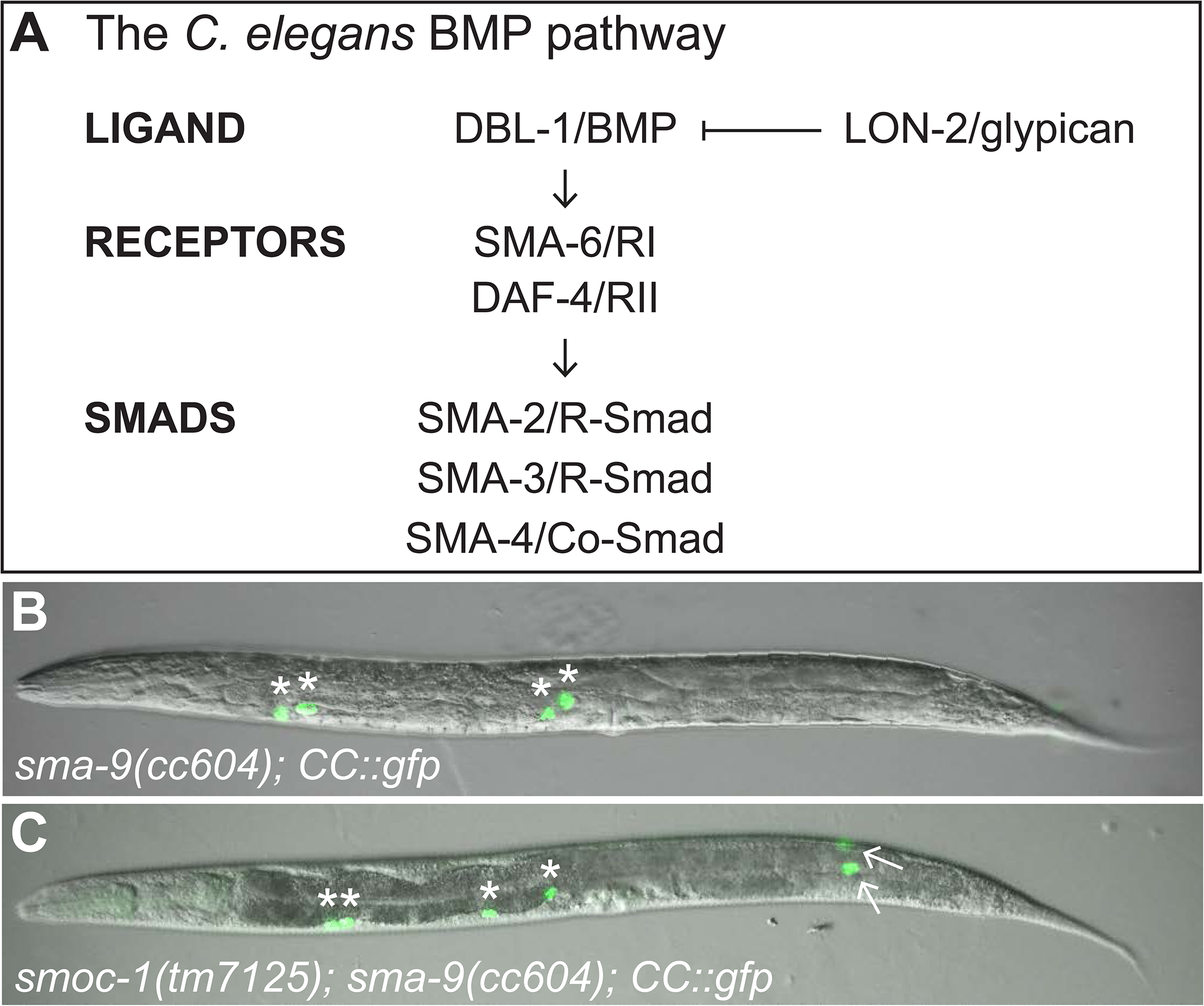
*smoc-1(0)* mutations suppress the *sma-9(0)* M lineage defect. (**A**) Schematic representation of the BMP signaling pathway in *C. elegans*. BMP: bone morphogenetic protein. RI: type I receptor. RII: type II receptor. R-Smad: receptor-associated Smad. Co-Smad: common mediator Smad. (**B-C**) Merged DIC and GFP images of L4 stage *sma-9(cc604)* (**B**) and *smoc-1(tm7125); sma-9(cc604)* (**C**) worms carrying the *CC::gfp* coelomocyte (CC) marker. Arrows indicate M-derived CCs. Asterisks (*) denote embryonically-derived CCs.

In this study, we report the identification and characterization of a new BMP modulator, which we have named SMOC-1. SMOC-1 is predicted to be a secreted protein that contains a thyroglobulin-like (TY) domain and an extracellular calcium-binding (EC) motif. We show here that SMOC-1 acts as a positive modulator of BMP signaling in *C. elegans*. We further demonstrate that SMOC-1 acts upstream of the ligand to regulate body size. We identified *smoc-1*-expressing cells, and demonstrated that SMOC-1 acts cell non-autonomously to regulate BMP signaling. Finally, we provide evidence that the function of SMOC proteins in the BMP pathway is conserved from worms to humans.

## MATERIALS & METHODS

### *C. elegans* strains

All strains were maintained at 20°C using standard culture conditions (BRENNER 1974) unless otherwise specified. Table 1 lists all the strains used in this study.

**Table 1.**
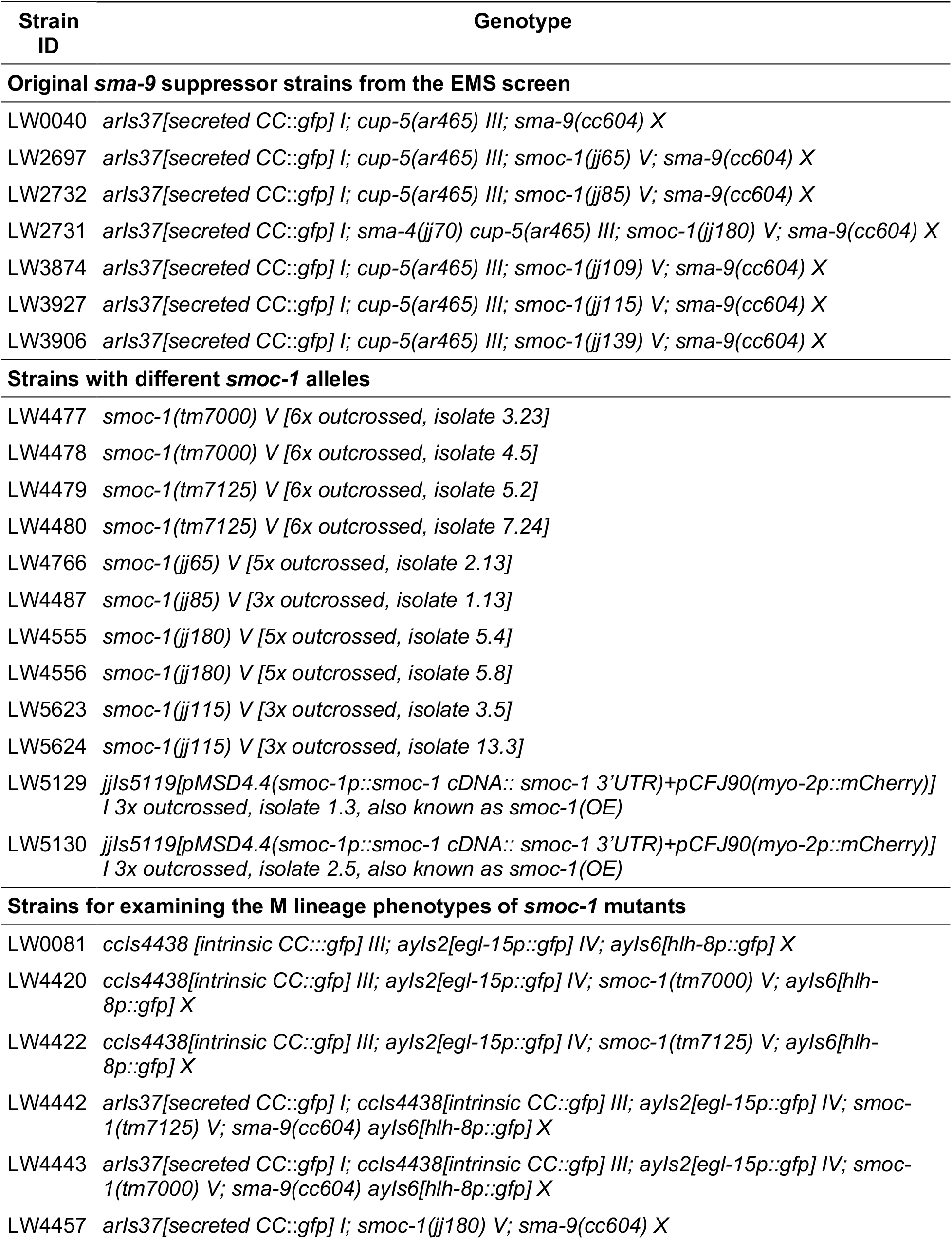
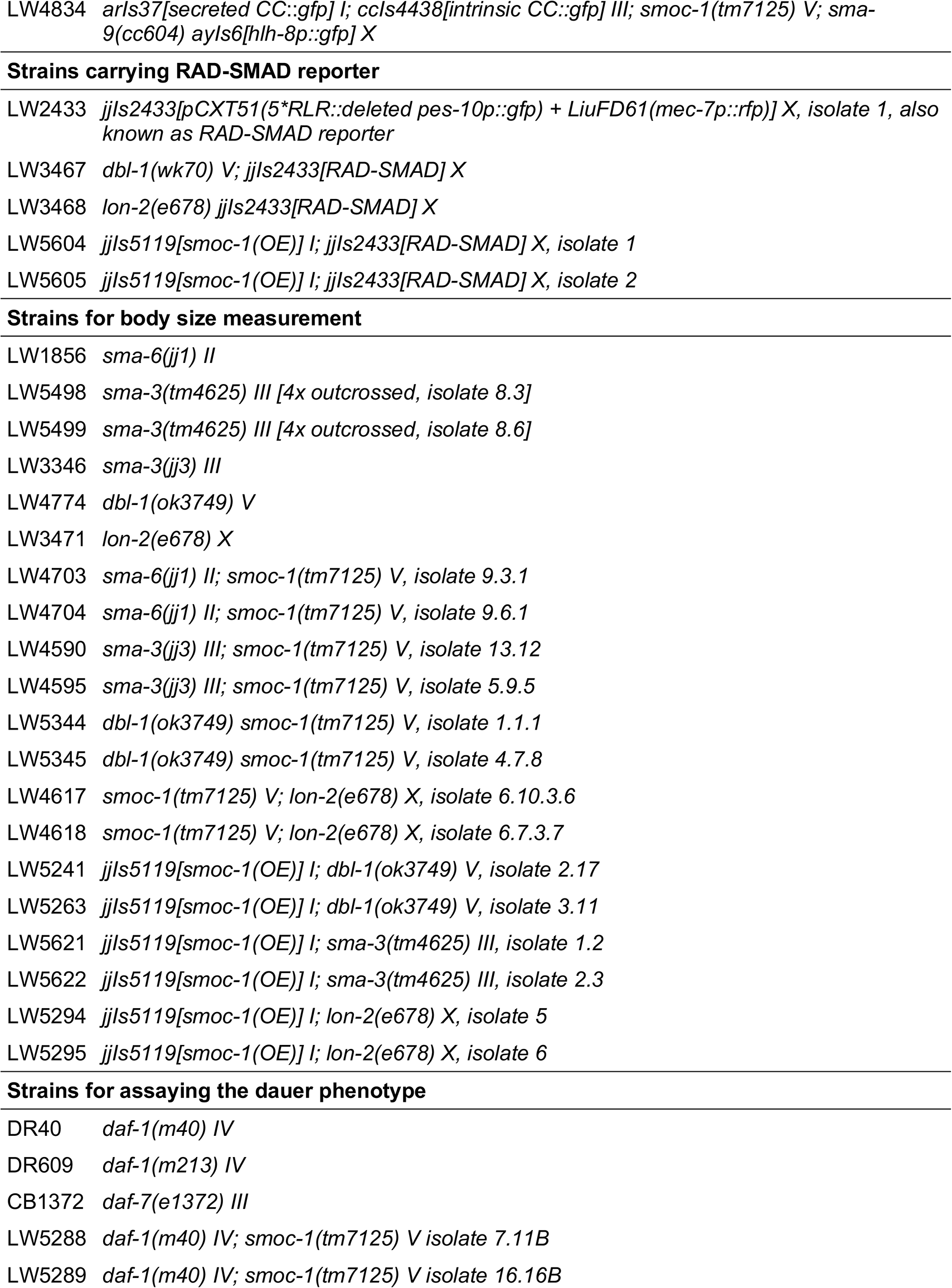
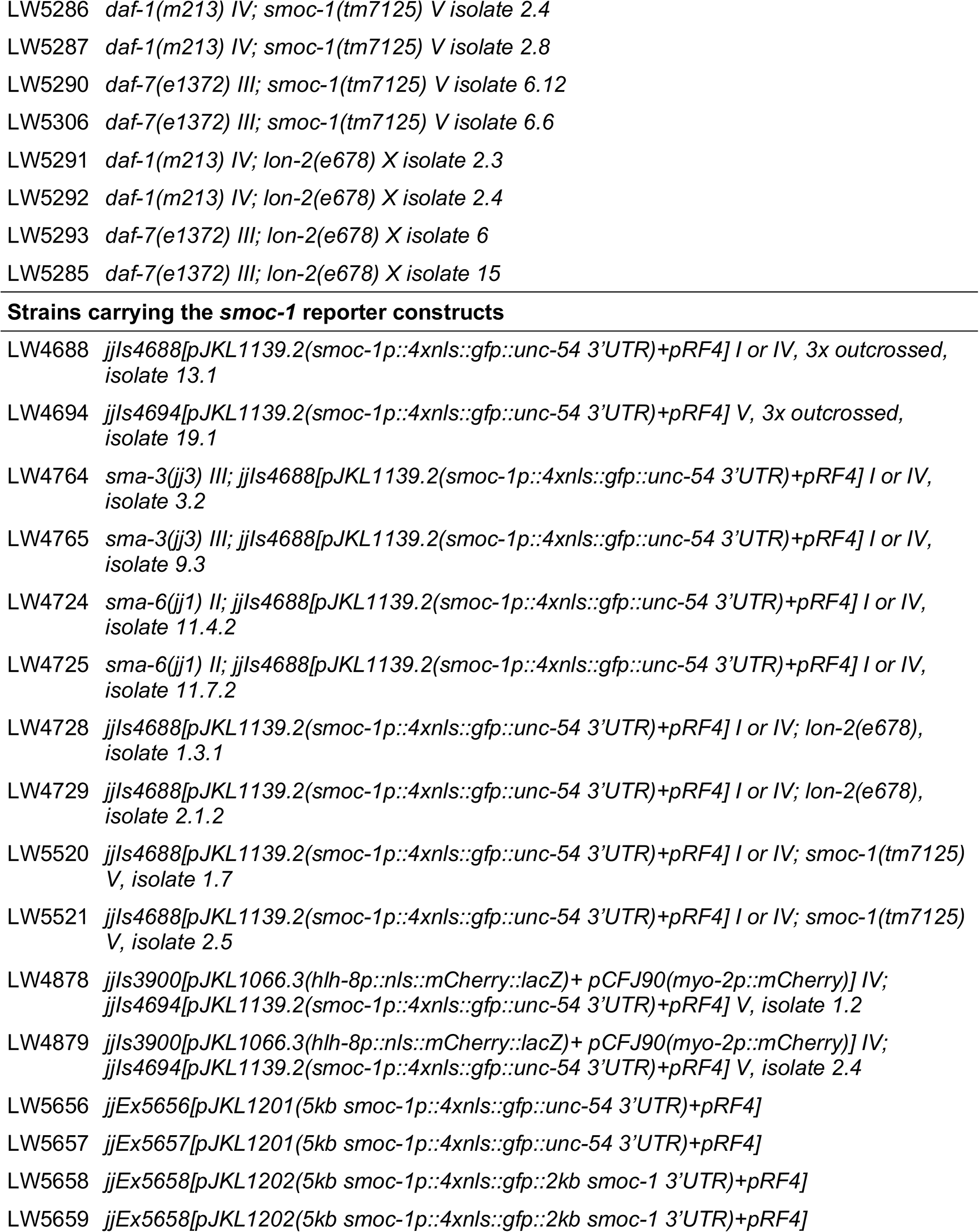
Strains used in this study.

### Plasmid constructs and transgenic lines

All plasmid constructs used in this study are listed in Table 2. The *smoc-1* open reading frame was amplified from the Vidal RNAi library (RUAL *et al.* 2004). Subsequent sequencing of the clone revealed the presence of a point mutation (S103P, Figure 2D), changing amino acid 103 from serine (TCC) to proline (CCC). Site directed mutagenesis was used to fix this point mutation. Plasmids containing the human *SMOC1* and *SMOC2* cDNAs were purchased from PlasmID, the DNA resource core at Harvard Medical School.

**Table 2.**
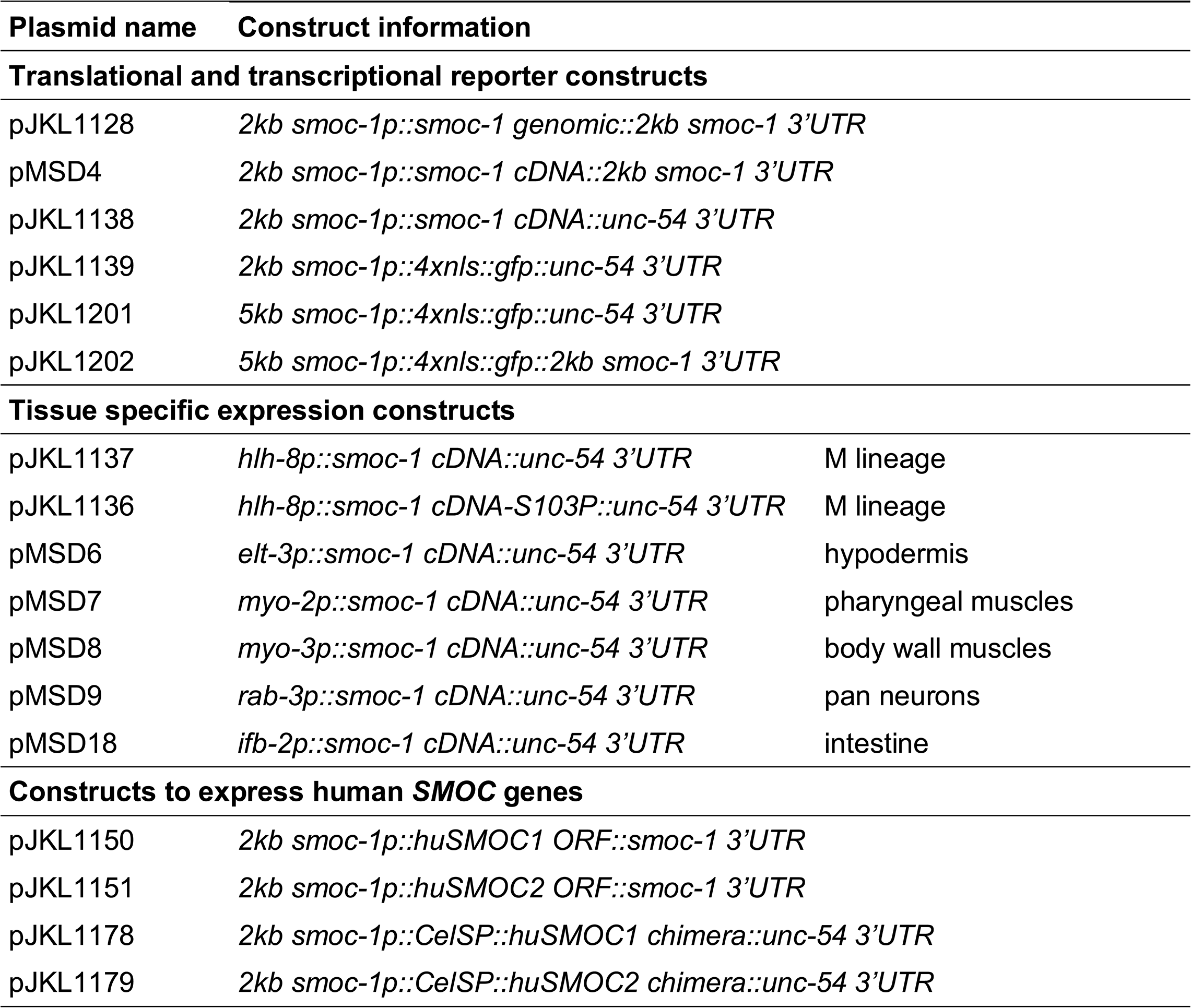
Plasmid constructs generated in this study.

**Fig 2.**
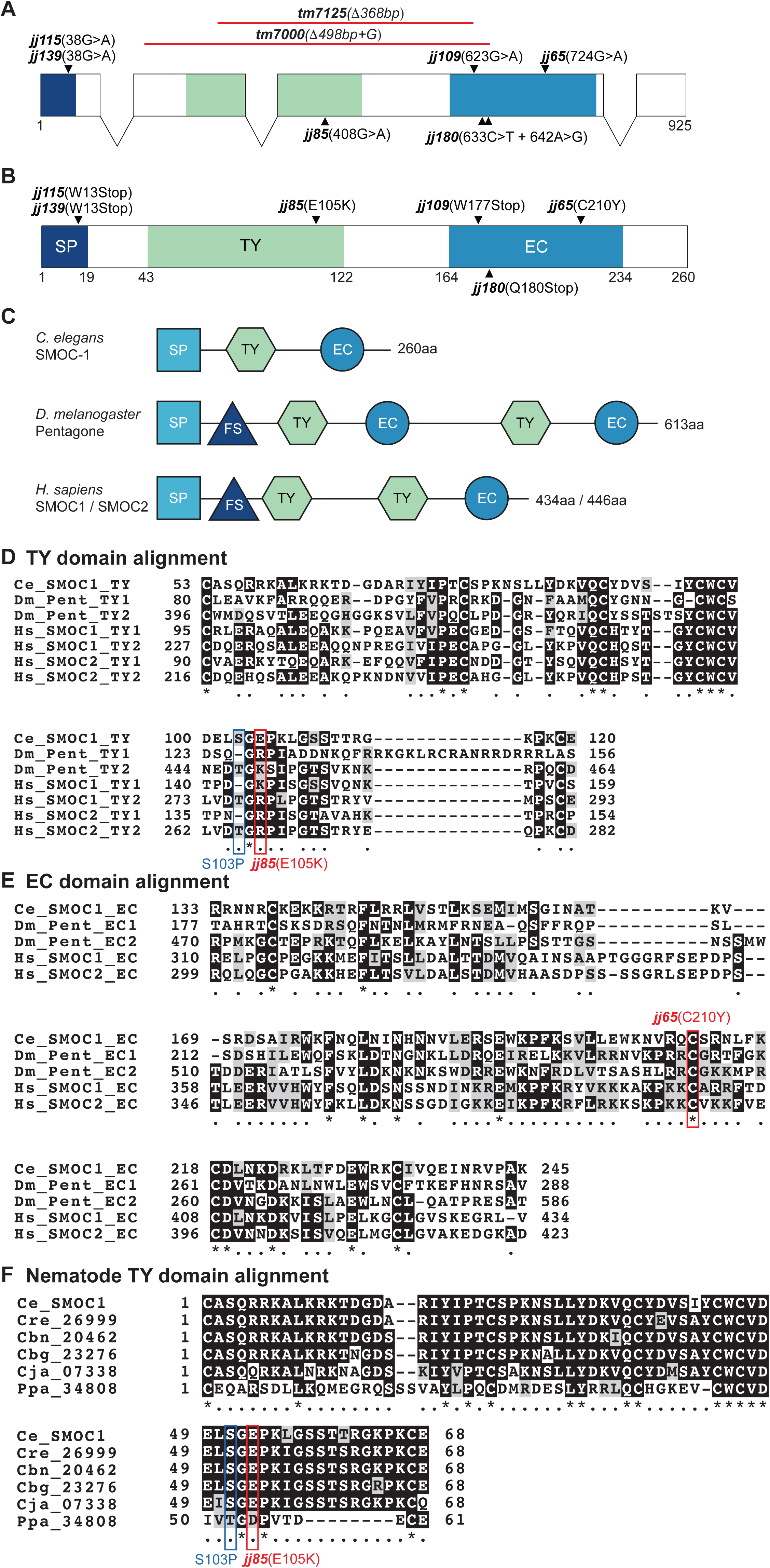
SMOC-1 is conserved from *C. elegans* to human. (**A-B**) Schematics of the *C. elegans smoc-1* gene (**A**) and the predicted SMOC-1 protein (B), respectively, showing the domain structure and the molecular lesions of various mutant alleles. SP: signal peptide. TY: thyroglobulin type I-like repeat. EC: secreted protein acidic and rich in cysteine (SPARC) extracellular calcium binding domain. (**C**) Schematic representation of *C. elegans* SMOC-1, *D. melanogaster* Pentagone, and *H. sapiens* SMOC1 and SMOC2, showing their domain structures. The two human SMOC proteins are of different lengths but share similar domain structures. FS: follistatin-like domain. (**D-E**) Alignment of the TY (**D**) and EC (**E**) domains from *C. elegans* SMOC-1, *D. melanogaster* Pentagone, and *H. sapiens* SMOC1 and SMOC2. Multiple copies of a certain domain in the same protein are numbered in order from the N-terminus to the C-terminus. Ce: *C. elegans*. Dm: *D. melanogaster*. Hs: *H. sapiens*. (**F**) Alignment of the TY domains from SMOC-1 homologs in various nematode species. Cel: *C. elegans*. Cre: *C. remanei*. Cbn: *C. brenneri*. Cbg: *C. briggsae*. Cja: *C. japonica*. Ppa: *Pristionchus pacificus*. In D-F, identical or conserved amino acids are shown on a black or grey background, respectively. Red boxes highlight residues mutated in certain *smoc-1* alleles. Blue box indicates the residue changed in a *smoc-1* cDNA clone that rendered the protein non-functional.

Transgenic strains were generated using the plasmid pRF4 *(rol-6(su1006))*, pCFJ90 (*myo-2p*::*mCherry*::*unc-54 3*’*UTR)*, or pJKL724 (*myo-3p::mCherry::unc-54 3’ UTR)* as a co-injection marker. Two transgenic lines with the best transmission efficiency were analyzed for each plasmid of interest. Integrated transgenic lines either overexpressing *smoc-1* (*jjIs5119*) or carrying the *smoc-1* transcriptional reporter (*jjIs4688* and *jjIs4694*) were generated using gamma-irradiation, followed by three rounds of outcrossing with N2 worms.

### Protein sequence alignment

Sequences where taken from Genbank (*C. elegans* SMOC-1 (T04F3.2), 179609; *C. remanei* CRE_26999, 9815068; *C. briggsae* CBG23276, 8578577; *D. melanogaster* Pent/Magu, 44850; *H. sapiens* SMOC1, 64093; *H. sapiens* SMOC2, 64094) or Wormbase (*C. brenneri* CBN20462; *C. japonica* CJA07338; *P. pacificus* PPA34808). TY and EC domains in SMOC proteins were predicted by Interpro (FINN *et al.* 2017). Domains were aligned using M-COFFEE Multiple Sequence Alignment (MSA) tool on the T-COFFEE server (version 11.00.d625267, (WALLACE *et al.* 2006)). ALN files were processed to produce alignment images using BOXSHADE.

### Microscopy

Epifluorescence and differential interference contrast (DIC) microscopy were conducted on a Leica DMRA2 compound microscope equipped with a Hamamatsu Orca-ER camera using the iVision software (Biovision Technology, Inc.). Subsequent image analysis was performed using Fiji (SCHINDELIN *et al.* 2012). RAD-SMAD reporter assay was carried out as previously described (TIAN *et al.* 2013).

### Body size measurements

Body size measurement assays were conducted as previously described (TIAN *et al.* 2013). Hermaphrodite worms were imaged at the L4.3 stage based on vulva development (MOK *et al.* 2015). Body sizes were measured from images using the segmented line tool of Fiji. An ANOVA and a Tukey HSD were conducted to test for differences in body size between genotypes using R (R CORE TEAM 2015).

### Suppression of *sma-9(0)* M-lineage defect (Susm) assay

For the Susm assay, worms were grown at 20°C and then the number of animals with 4 CCs and 6 CCs were tallied across three to seven plates for each genotype. For the Susm rescue experiments, we generated general linear models (GLMs) with binomial errors, and a logit link function designating transgene as the explanatory function to test for differences between transgenic and non-transgenic groups within a line.

### Dauer formation assay

Dauer formation assay was conducted under non-dauer-inducing conditions as previously described (VOWELS AND THOMAS 1992). Ten adult hermaphrodites were placed on a six centimeter NGM plate (five plates per strain at each temperature) and allowed to lay eggs for less than eight hours. Adults were removed and plates were placed at the test temperature. When non-dauer worms became young adults, the numbers of dauer and non-dauer worms on each plate were scored. Using R, we tested for differences in dauer formation between genotypes using an ANOVA followed by a TukeyHSD.

### Data availability statement

Strains and plasmids are available upon request. The authors affirm that all data necessary for confirming the conclusions of the article are present within the article, figures, and tables.

## RESULTS

### Mutations in T04F3.2 suppress the mesoderm defects of *sma-9(0)* mutants

In a previous *sma-9* suppressor screen, we uncovered a novel complementation group named *susm-1* that includes three alleles, *jj65, jj85* and *jj180* (LIU *et al.* 2015) (Table 3), which suppressed the *sma-9(0)* M lineage defect at high penetrance. Whole genome sequencing (WGS) of the three alleles identified molecular lesions in the uncharacterized gene T04F3.2: *jj65* and *jj85* are missense mutations C210Y and E105K, respectively, while *jj180* is a nonsense mutation Q180Stop (Figure 2A,B). To confirm that T04F3.2 is the corresponding gene for this complementation group, we obtained two deletion alleles that delete most of the coding region of T04F3.2, *tm7000* and *tm7125* (Figure 2A), and found that both alleles suppressed the *sma-9(0)* M lineage defect to near 100% (Table 3, Figure 1C). Pairwise complementation tests between *tm7000* and *jj65, jj85* or *jj180*, showed that *tm7000* failed to complement all three alleles in their suppression of the *sma-9(0)* M lineage defect (Table 3). Subsequent *sma-9(0)* suppressor screens conducted in the lab identified three additional alleles of this complementation group, *jj109, jj115,* and *jj139*. WGS followed by Sanger sequencing showed that all three alleles contain nonsense mutations in T04F3.2: W13Stop for both *jj115* and *jj139*, and W176Stop for *jj109* (Figure 2A,B). Finally, a transgene containing the T04F3.2 genomic region including 2kb upstream sequences, the entire coding region with introns, and 2kb downstream sequences rescued the *sma-9(0)* suppression phenotype of *tm7125* mutants (Table 3). Collectively, these results demonstrated that T04F3.2 is the corresponding gene for the *susm-1* locus. The nature of the molecular lesions in *tm7000, tm7125, jj109, jj115, jj139,* and *jj180*, the near 100% penetrance of their Susm phenotypes, and their similar body size phenotypes (see below), suggest that all of these alleles are null alleles. For ease of genotyping, most of our subsequent analysis was carried out using the *tm7125* allele.

**Table 3.**
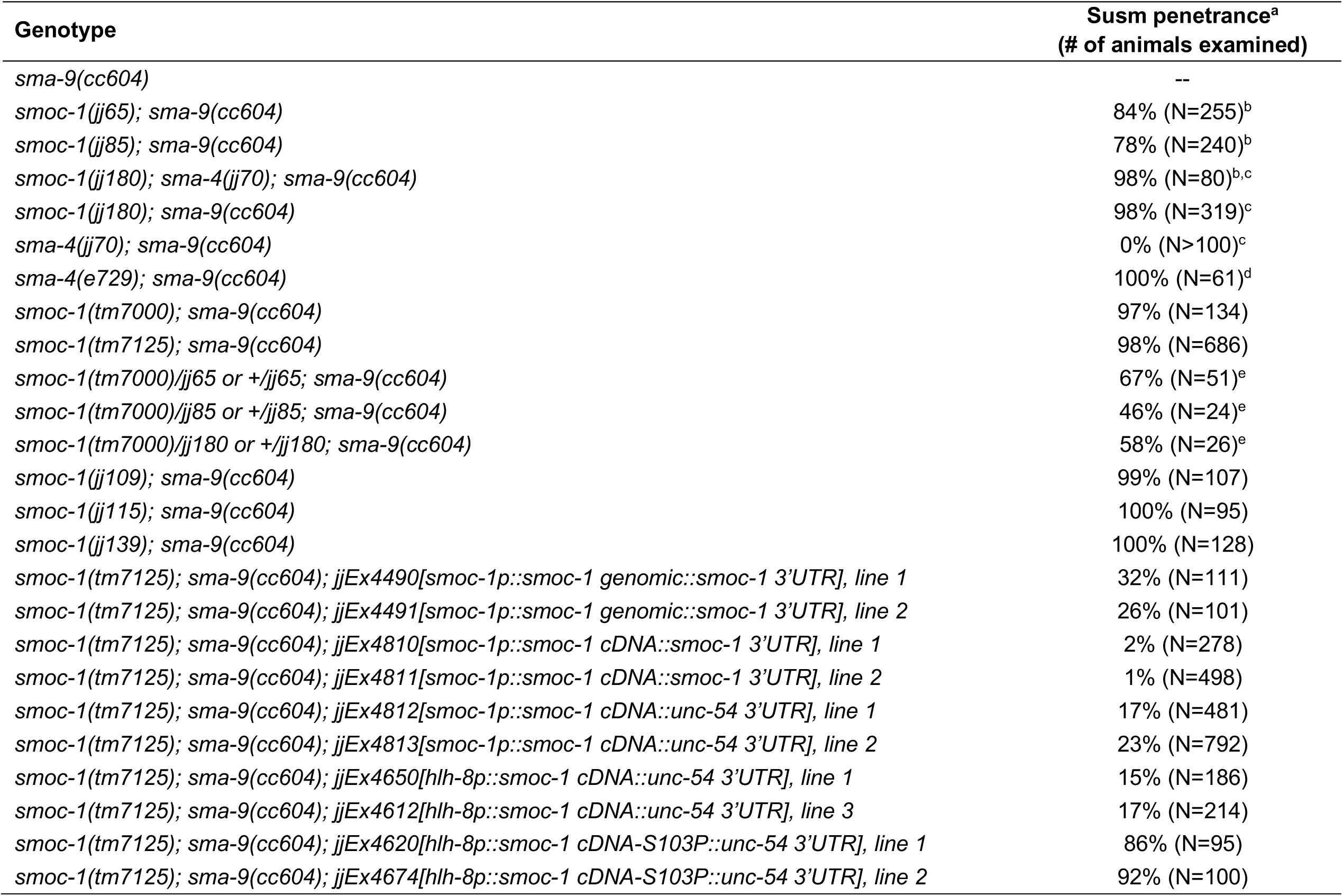
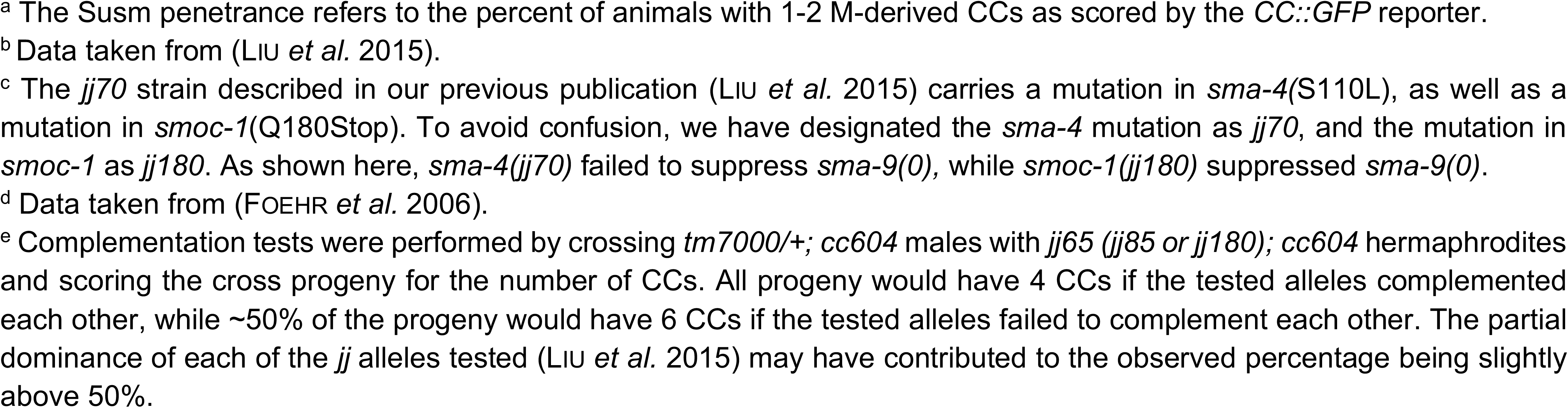
Mutations in *smoc-1* suppress the *sma-9(0)* M lineage defects.

### T04F3.2 encodes a predicted Secreted MOdular Calcium-binding protein SMOC-1

T04F3.2 is predicted to encode a protein of 260 amino acids. It contains a predicted signal peptide (SP), a thyroglobulin type I-like repeat (TY), and a secreted protein acidic and rich in cysteine (SPARC) extracellular calcium (EC) binding region (Figure 2). The EC domain is predicted to contain a pair of helix-loop-helix EF hand calcium-binding motifs (HOHENESTER *et al.* 1996; VANNAHME *et al.* 2002). The predicted T04F3.2 protein is most similar to the human secreted modular calcium-binding proteins SMOC1 and SMOC2 (VANNAHME *et al.* 2002; VANNAHME *et al.* 2003), and the *Drosophila melanogaster* SMOC homolog Pentagone/Magu (VUILLEUMIER *et al.* 2010). A BLAST search against the *C. elegans* genome showed that T04F3.2 is the only SMOC homolog. We have, therefore, named this gene *smoc-1* and its corresponding protein SMOC-1.

SMOC proteins are matricellular proteins that are in the same family as SPARC/BM-40/osteonectin (BRADSHAW 2012). The domain arrangement of SMOC proteins varies across species. The *C. elegans* SMOC-1 protein is predicted to have one TY domain, one EC domain, and completely lack the follistatin (FS) domain that is present in other SMOC proteins (Figure 2C). Within the TY domain, SMOC-1 shares about 30% amino acid identity and 50% similarity with human SMOC1 and SMOC2, and contains a CWCV tetrapeptide sequence and an additional four conserved cysteines that are characteristic of the TY domain (Figure 2D). The EC domain of SMOC-1 shares about 25% amino acid identity and 45% similarity with those of the human SMOC proteins. Among the conserved residues in the EC domain are four cysteines thought to be involved in disulfide bond formation (BUSCH *et al.* 2000).

The locations of the molecular lesions in our *smoc-1* mutant alleles suggest that both the TY domain and the EC domain are important for SMOC-1 function. *jj85* is a mutation in the TY domain, changing amino acid 105 from a glutamic acid to a lysine (E105K, Figure 2B,D). Although the change appears to make this residue more similar to its counterpart (arginine or lysine) in the fly and human SMOC proteins (Figure 2D), we noted that E105 is conserved in multiple nematode species (Figure 2F). We also obtained a *smoc-1* cDNA clone that has a single base mutation changing amino acid 103 from a conserved serine to proline (Figure 2D). This mutant *smoc-1* cDNA (S103P) failed to rescue the *smoc-1(0)* Susm phenotype, while the wild-type (WT) *smoc-1* cDNA under the same regulatory elements successfully rescued the *smoc-1(0)* Susm phenotype (Table 3), again highlighting the importance of the TY domain for SMOC-1 function. Similarly, the EC domain is also critical for SMOC-1 function, because a change of the conserved cysteine residue at amino acid 210 to tyrosine (C210Y) in *jj65* significantly compromised the function of SMOC-1 (Figure 2B,E, Table 3).

### SMOC-1 functions within the BMP pathway to positively regulate BMP signaling

We have previously shown that mutations in BMP pathway components specifically suppress the *sma-9(0)* M lineage defect (FOEHR *et al.* 2006; LIU *et al.* 2015). The highly penetrant Susm phenotype of multiple *smoc-1* alleles suggests that SMOC-1 may function in the BMP pathway. BMP pathway mutants are known to exhibit altered body sizes (SAVAGE-DUNN AND PADGETT 2017). We measured the body sizes of *smoc-1* single mutant animals and found that they all have a reproducibly smaller body size (∼95%) compared to WT animals at the same developmental stage (Figure 3A,B,D). This smaller body size can be rescued by a WT *smoc-1* transgene (Figure 3D). Moreover, transgenic *smoc-1* mutant animals carrying this transgene are significantly longer than WT animals (Figure 3D). This is likely due to the repetitive nature of the transgene generated using standard *C. elegans* transgenic approaches, which often results in over-expression of the transgene (MELLO *et al.* 1991). We have subsequently integrated the WT *smoc-1* transgene in the WT background (*jjIs5119*, Table 1). Again, *jjIs5119* (which we have referred to as *smoc-1(OE)*) animals are significantly longer than WT animals (Figure 4B). Thus, *smoc-1* appears to function in a dose-dependent manner to positively regulate body size.

**Fig 3.**
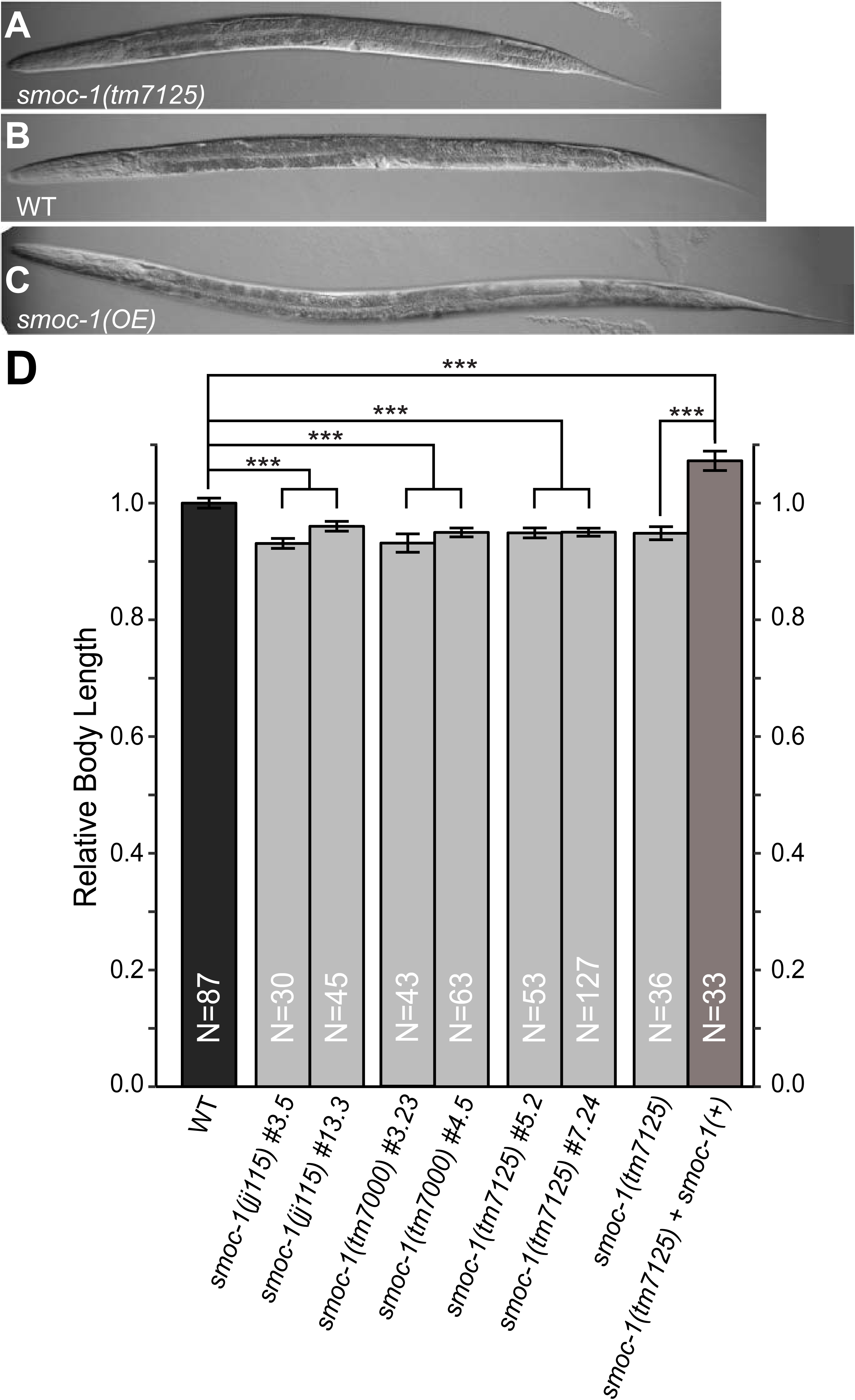
SMOC-1 regulates body size. (**A-C**) DIC images showing *smoc-1(tm7125)* (**A**), WT (**B**), and *smoc-1(OE)* (**C**) worms at the larval L4.3 stage. (**D**) Relative body length of developmental stage-matched WT and various *smoc-1* mutant worms. Each *smoc-1* mutant allele was outcrossed with N2 for at least three times, and two independent isolates for each allele (#s following the allele name) were used for body size measurement. The *smoc-1(+)* transgene was pMSD4*[2kb smoc-1p::smoc-1 cDNA::2kb smoc-1 3’UTR]*. The body length of WT worms was set to 1.0. Error bars represent 95% confidence interval (CI). An ANOVA followed by TukeyHSD was used to test for differences between genotypes. *** *P*<0.0001.

**Fig 4.**
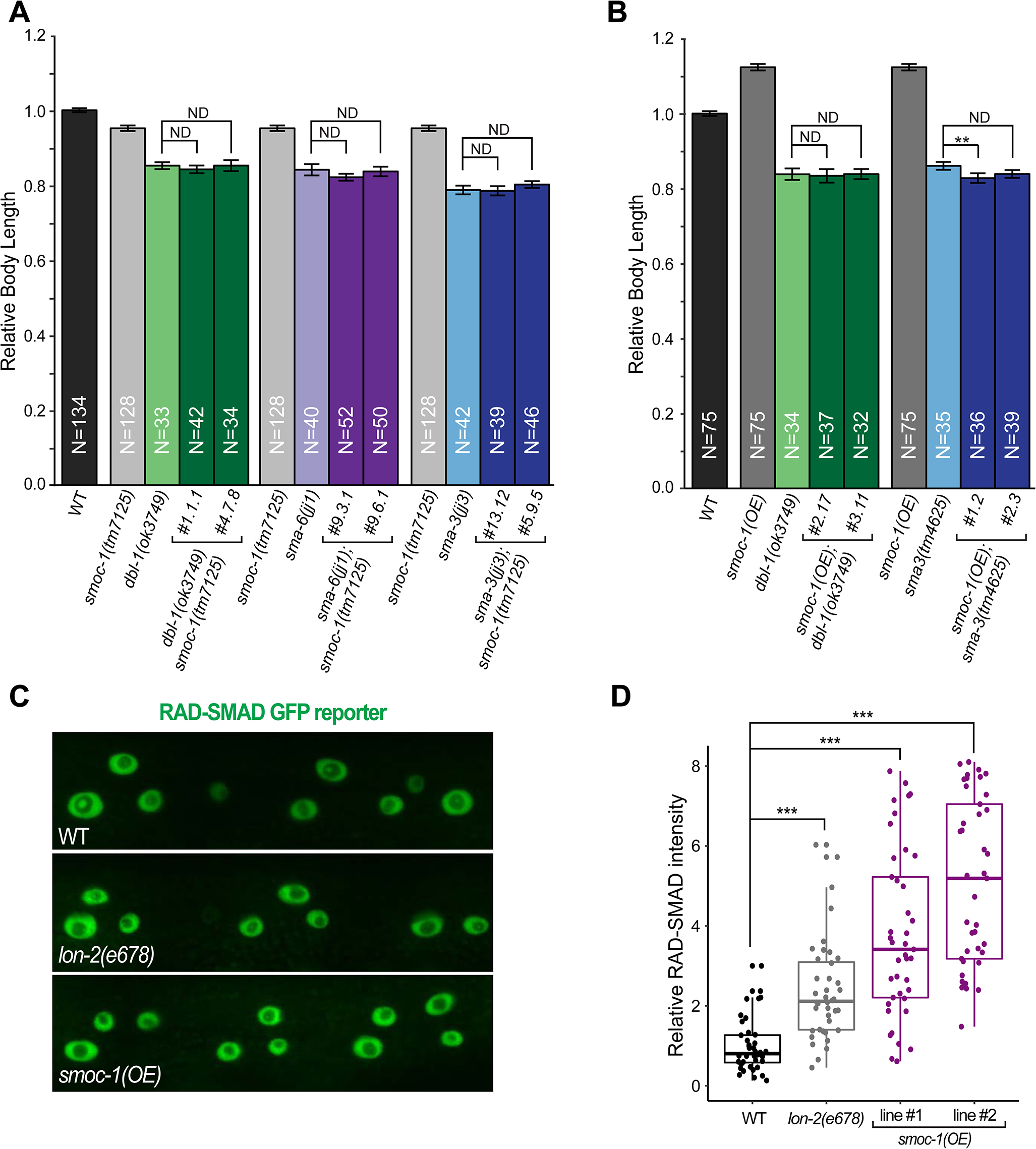
SMOC-1 functions through the BMP ligand to positively regulate BMP signaling. (**A-B**) Relative body length of developmental stage-matched WT and various mutant worms, including double mutants between *smoc-1(tm7125)* and null mutants in the BMP pathway (**A**), and double mutants between *smoc-1(OE)* and null mutants in the BMP pathway (**B**). Two independent isolates for each double mutant combination were used for body size measurement. The body length of WT worms was set to 1.0. Error bars represent 95% CI. (**C**) Representative GFP images showing RAD-SMAD reporter expression in hypodermal nuclei of WT, *lon-2(e678)*, and *smoc-1(OE)* worms, respectively. (**D**) Boxplot showing the relative RAD-SMAD GFP fluorescence intensity in WT (set to 1.0), *lon-2(e678)*, and two independent isolates of *smoc-1(OE)* worms. Each data point represents an average of the GFP fluorescence intensity from five hypodermal nuclei in one worm. Approximately 40 worms were examined per genotype. For panels A, B and D, an ANOVA followed by TukeyHSD was used to test for differences between genotypes. ND: no difference. *** *P*<0.0001.

To determine whether *smoc-1* functions within the BMP pathway to regulate body size, we generated double mutants between *smoc-1(tm7125)* and null mutations in various BMP pathway components, and measured their body lengths. As shown in Figure 4A, *dbl-1(ok3749) smoc-1(tm7125)* double mutants were as small as *dbl-1(ok3749)* single mutants. Similarly, *sma-3(jj3); smoc-1(tm7125)* and *sma-6(jj1); smoc-1(tm7125)* double mutants were as small as *sma-3(jj3)* and *sma-6(jj1)* single mutants, respectively. These observations indicate that *smoc-1* functions within the BMP pathway, rather than in a parallel pathway, to regulate body size.

In addition to body size, BMP pathway mutants also exhibit male tail defects and the mutant males cannot mate (SAVAGE *et al.* 1996; KRISHNA *et al.* 1999; SUZUKI *et al.* 1999). We generated *smoc-1(tm7125)* males and found that they mated well with WT hermaphrodites to produce cross progeny, suggesting that *smoc-1(tm7125)* males do not have severe male tail patterning defects. This is not surprising as previous studies have demonstrated that male tail development is not affected when there is a partial reduction of BMP signaling (KRISHNA *et al.* 1999).

We also examined the expression of the RAD-SMAD reporter, which we have previously shown to serve as a direct readout of BMP signaling (TIAN *et al.* 2010). While *smoc-1* null mutants did not exhibit significant changes in the expression of the RAD-SMAD reporter (data now shown), the *smoc-1(OE)* lines showed a significant increase in the level of RAD-SMAD reporter expression (Figure 4C,D). We reasoned that the change of RAD-SMAD reporter expression in *smoc-1(0)* mutants may be too small to detect given that *smoc-1(0)* mutants only exhibit about 5% reduction in body size compared to WT animals (see above). Nevertheless, our findings are consistent with SMOC-1 acting in the BMP pathway to positively promote BMP signaling.

### SMOC-1 functions through the BMP ligand to promote BMP signaling in regulating body size

The long body size phenotype caused by *smoc-1* overexpression provided us with a useful tool to determine where in the BMP signaling pathway SMOC-1 functions. We conducted genetic epistasis analysis by generating double mutants between *smoc-1(OE)* and null mutations in core components of the BMP pathway that are known to cause a small body size. As shown in Figure 4B, *smoc-1(OE); dbl-1(ok3749)* double mutants and *smoc-1(OE); sma-3(tm4625)* double mutants are as small as *dbl-1(ok3749)* and *sma-3(tm4625)* single mutants, respectively. These results provide further support to the conclusion that SMOC-1 functions within the BMP pathway to regulate body size. More importantly, our genetic epistasis results demonstrate that SMOC-1 functions upstream of and is dependent on the function of the BMP ligand DBL-1 to regulate body size.

### SMOC-1 antagonizes the function of LON-2/glypican to modulate BMP signaling in regulating body size

Previous studies have shown that the glypican LON-2 functions upstream of DBL-1/BMP and acts as a negative regulator of BMP signaling (GUMIENNY *et al.* 2007). We performed double mutant analysis and dissected the relationship between SMOC-1 and LON-2/glypican. We first measured the body length of double null mutants between *smoc-*1 and *lon-2*. As shown in Figure 5A, *smoc-1(tm7125); lon-2(e678)* double null mutants exhibited an intermediate body size compared to either single null mutant. In particular, the body size of *smoc-1(tm7125); lon-2(e678)* double mutants is similar to that of WT animals. These observations suggest that SMOC-1 and LON-2/glypican antagonize each other in regulating body size. Interestingly, *smoc-1(OE); lon-2(e678)* worms are longer than either *smoc-1(OE)* animals or *lon-2(e678)* single mutants (Figure 5B). Thus, over-expressing *smoc-1* is capable of further increasing the body size of worms that completely lack LON-2/glypican. Taken together, our genetic analysis between *lon-2* and *smoc-1* suggests that SMOC-1 antagonizes the function of LON-2/glypican in regulating body size, and that SMOC-1 also has LON-2/glypican-independent function(s) in promoting BMP signaling.

**Fig 5.**
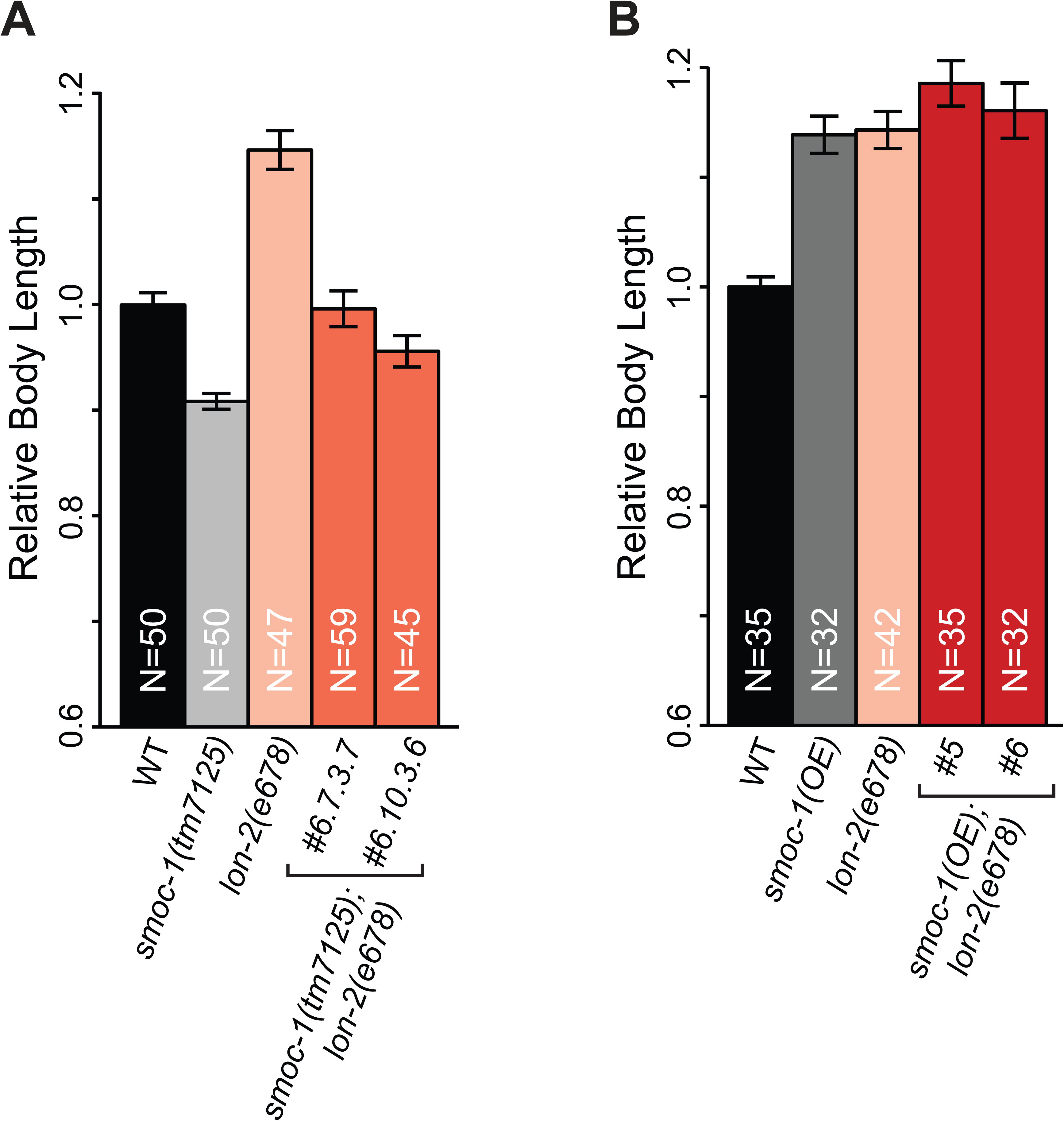
SMOC-1 antagonizes LON-2/glypican in regulating body size. Relative body length of developmental stage-matched WT (set to 1.0) and various mutant worms, including double mutants between *smoc-1(tm7125)* null and *lon-2(e678)* null (**A**), and double mutants between *smoc-1(OE)* and *lon-2(e678)* null (**B**). The body size of *smoc-1(tm7125)*; *lon-2(e678)* double mutants is similar to that of WT animals, while *smoc-1(OE); lon-2(e678)* double mutants are longer than either one. Error bars represent 95% CI. An ANOVA followed by TukeyHSD was used to test for differences between genotypes. ND: no difference. * *P*<0.01. *\*\* P*<0.001. *\*\*\* P*<0.0001.

### SMOC-1 does not play a major role in the TGFβ-like dauer pathway

In addition to the BMP pathway, *C. elegans* has a TGFβ-like signaling pathway that regulates dauer development (SAVAGE-DUNN AND PADGETT 2017). To determine if SMOC-1 plays a role in the TGFβ-like dauer pathway, we first assayed dauer formation of worms with different levels of *smoc-1* expression. *smoc-1(tm7125)* and *smoc-1(OE)* single mutant worms did not exhibit any constitutive or defective dauer formation phenotype at any of the temperatures tested (Table 4, data not shown), suggesting that SMOC-1 does not play a major role in the TGFβ-like dauer pathway. Next, we generated double mutant worms carrying both *smoc-1(tm7125)* and mutations in the TGFβ ligand DAF-7/TGFβ or the type 1 receptor DAF-1/RI (GEORGI *et al.* 1990; REN *et al.* 1996), and examined them for the constitutive dauer formation (Daf-c) phenotype (Table 1). While *smoc-1(tm7125)* partially suppressed the Daf-c phenotype of *daf-7(e1372)* at 20°C, a similar trend was not observed at either 15°C or at 25°C. Similarly, *smoc-1(tm7125)* did not exhibit any consistent suppression or enhancement of the Daf-c phenotype of two *daf-1* mutant alleles (Table 4). These results suggest that SMOC-1 does not play a major role in the TGFβ-like dauer pathway, although we cannot rule out a minor buffering function of SMOC-1 in this pathway.

**Table 4.**
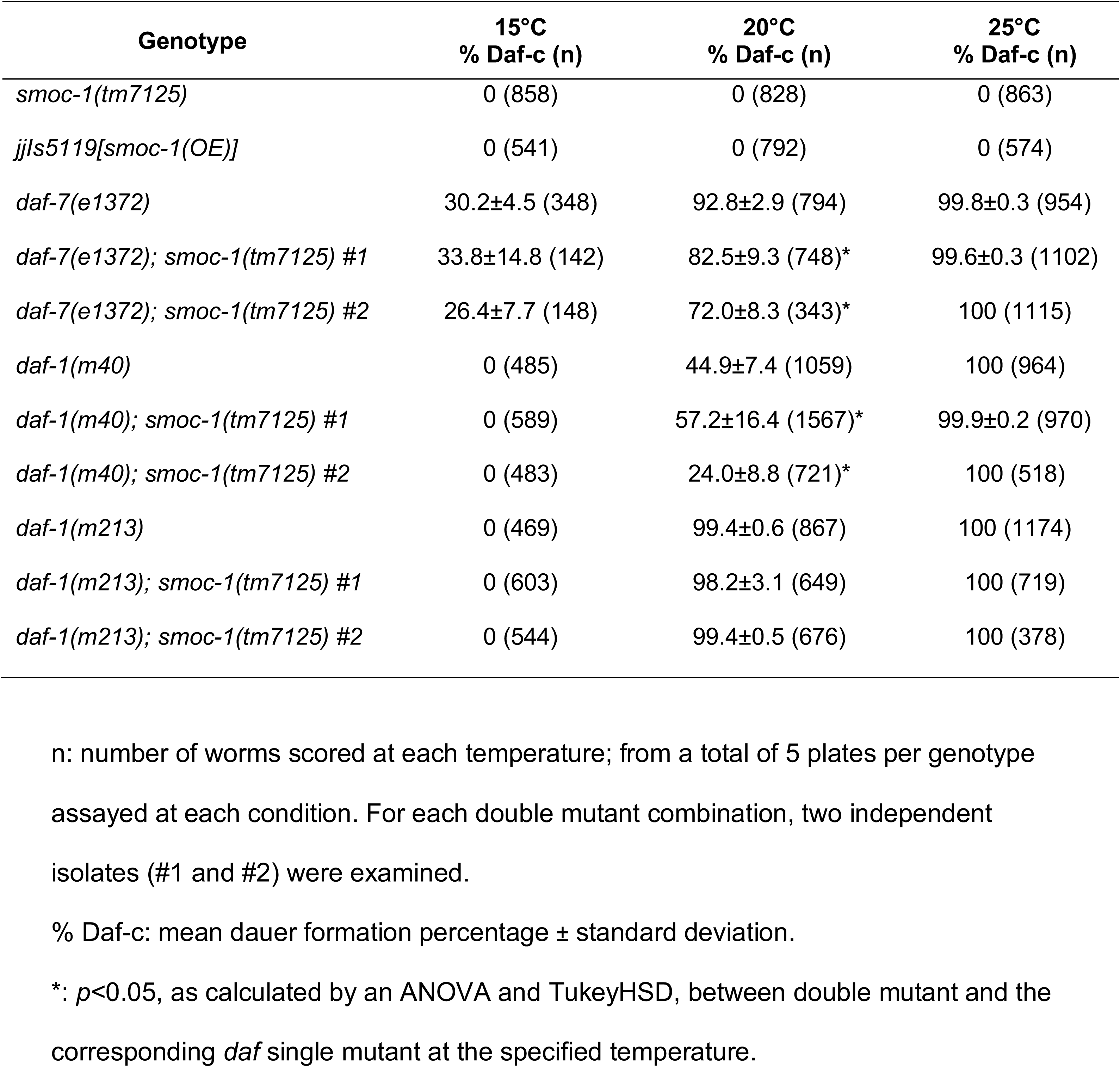
SMOC-1 does not play a significant role in the TGFβ dauer pathway.

Because of the genetic interaction that we observed between *smoc-1* and *lon-2*, we also tested whether LON-2/glypican plays a role in the TGFβ-like dauer pathway by performing similar double mutant analysis as described for *smoc-1*. At 20°C, *lon-2(e678)* showed partial suppression of the Daf-c phenotype of *daf-7(e1372)* (Table 5), but a similar trend was not observed at 15°C or at 25°C (Table 5). As seen with *smoc-1(tm7125), lon-2(e678)* also did not consistently enhance or suppress the Daf-c phenotype of a TGFβ receptor mutation, *daf-1(m213)*. Thus, like SMOC-1, LON-2 does not appear to play a major role, but may play a minor modulatory role, in the TGFβ dauer pathway.

**Table 5.** LON-2 does not play a significant role in the TGFβ dauer pathway.

| **Genotype** | **15°C**<br>**% Daf-c (n)** | **20°C**<br>**% Daf-c (n)** | **25°C**<br>**% Daf-c (n)** |
| --- | --- | --- | --- |
| *lon-2(e678)* | 0 (807) | 0 (799) | 0 (1090) |
| *daf-7(e1372)* | 22.7±32.1 (141) | 83.3±6.5 (257) | 100 (357) |
| *daf-7(e1372); lon-2(e678)* #1 | 25.3±8.7 (435) | 59.4±16.4 (239)* | 100 (749) |
| *daf-7(e1372); lon-2(e678)* #2 | 31.7±4.8 (249) | 66.7±11.7 (426) | 100 (840) |
| *daf-1(m213)* | 0.3±1.0 (313) | 97.9±26.0 (570) | 100 (853) |
| *daf-1(m213); lon-2(e678)* #1 | 0.2±0.3 (575) | 92.1±2.9 (643) | 100 (1149) |
| *daf-1(m213); lon-2(e678)* #2 | 0.5±0.8 (654) | 77.7±5.0 (515) | 100 (832) |

n: number of worms scored at each temperature; from a total of 5 plates per genotype assayed at each condition. For each double mutant combination, two independent isolates (#1 and #2) were examined.
% Daf-c: mean dauer formation percentage ± standard deviation.
\*: $p < 0.05$ , as calculated by an ANOVA and TukeyHSD, between double mutant and the corresponding *daf* single mutant at the specified temperature.

### *smoc-1* is expressed in the pharynx, intestine, and posterior hypodermis

Since *smoc-1* is predicted to encode a secreted protein, we first attempted to identify the cells that express *smoc-1*. As described above, a *smoc-1* genomic fragment containing 2kb upstream sequences, the entire coding region with introns, and 2kb downstream sequences (pJKL1128, Table 2) can rescue the Susm and body size phenotypes of *smoc-1(0)* mutants (Figure 6A, Table 3). The same promoter element driving the *smoc-1* cDNA with its own 3’UTR or with the *unc-54* 3’UTR rescued both the small body size and the Susm phenotypes of *smoc-1(0)* mutants (Figure 6A; Table 3), suggesting that the regulatory elements required for SMOC-1 function in BMP signaling reside in the 2kb upstream sequences. We therefore generated a transcriptional reporter *pJKL1139[smoc-1 2kb promoter::4xnls::gfp::unc-54 3’UTR]* (Table 2). We also generated two additional transcriptional reporters using 5kb *smoc-1* upstream sequences (*pJKL1201[smoc-1 5kb promoter::4xnls::gfp::unc-54 3’UTR]* and *pJKL1202[smoc-1 5kb promoter::4xnls::gfp::2kb smoc-1 3’UTR]*, Table 2). All three reporters showed similar expression patterns in transgenic animals. We therefore focused on *pJKL1139[smoc-1 2kb promoter::4xnls::gfp::unc-54 3’UTR]* and generated integrated transgenic lines carrying this reporter (*jjIs4688* and *jjIs4694*, Table 1) for further analysis.

**Fig 6.**
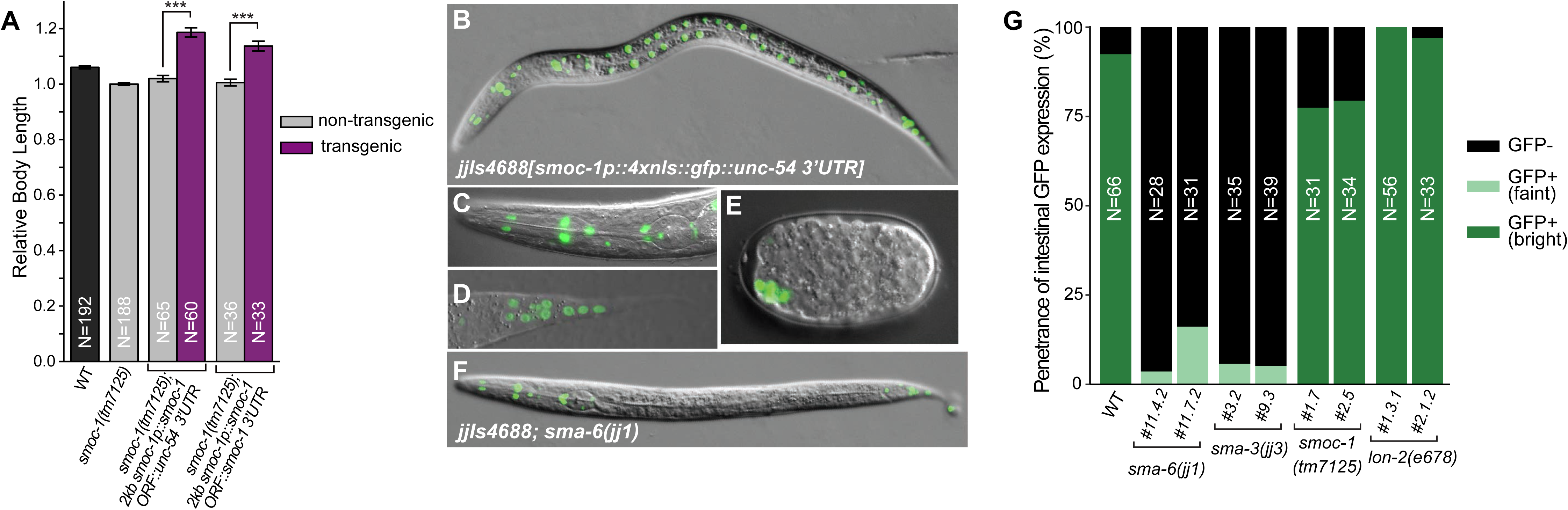
*smoc-1* is expressed in multiple tissues and its intestinal expression is positively regulated by BMP signaling. (**A**) Expression of *smoc-1* cDNA under different regulatory elements to test for rescue of the body size phenotype of *smoc-1(tm7125)* worms. For each construct, two independent transgenic lines were examined and the data were combined and averaged. Body sizes are relative to *smoc-1(tm7125)* mutant worms (set to 1.0), and all measurements were done on the same day. Error bars represent 95% CI. *** *P*<0.0001. (**B-F**) Merged GFP and DIC images of wildtype worms (**B-E**) and a *sma-6(jj1)* mutant worm (**F**) carrying the integrated *smoc-1* transcriptional reporter *jjIs4688* (Table 1). GFP expression is detectable a bean stage embryo (**E**), and in cells of the pharynx (**B-C**), intestine (**B**), and posterior hypodermis (**B, D**) in a WT larva. GFP expression in the intestine, but not in the pharynx or posterior hypodermis, is significantly reduced in *sma-6(jj1)* (**F**). Images are side views with anterior to the left and dorsal up. (**G**) Quantification of the penetrance of L4 stage animals showing intestinal expression of the *smoc-1* transcriptional reporter in wildtype and various BMP pathway mutants. Two independent isolates were assessed for each gene tested.

The integrated *smoc-1* transcriptional reporter showed strong GFP expression. GFP was first detectable in several cells located in the anterior of bean stage embryos (Fig 6F). In the developing larvae, GFP is expressed in cells of the pharynx, the intestine and the posterior hypodermis (Fig 6B). Pharyngeal cells expressing *smoc-1p::gfp* include the epithelial cells e2, the marginal cells mc1 and mc2, the M4 neuron, and all six of the pharyngeal/intestinal valve cells (Figure 6C). Cells of the posterior hypodermis expressing *smoc-1p::gfp* include hyp8, hyp9, hyp10, and hyp11 (Fig 6D). Expression in these tissues persisted from the L1 larval stage through adulthood. We noted that while all transgenic animals showed GFP expression in the pharynx and the posterior hypodermis, a small fraction of animals (∼8%) did not exhibit GFP expression in all or some of the intestinal cells (Figure 6G). We observed no GFP expression in any other tissues, including the nerve cord, body wall muscles (BWMs), or the M lineage. Thus, *smoc-1* is expressed in cells of the pharynx, intestine, and posterior hypodermis.

### Intestinal expression of *smoc-1* is positively regulated by BMP signaling

We next asked whether *smoc-1* expression is regulated by the BMP pathway or by SMOC-1 itself. We introduced the integrated *smoc-1* transgenic reporter into BMP pathway null mutants, including *sma-3(jj3), sma-6(jj1), lon-2(e678)* and *smoc-1(tm7125)* mutants (Table 1), and examined the expression pattern of the GFP reporter. Intriguingly, while the expression pattern and expression level of the GFP reporter in the pharynx and posterior hypodermis remained relatively constant in all mutant background examined, in *sma-6(jj1)* and *sma-3(jj3)* mutants there was a significant decrease in the percentage of animals that exhibited GFP expression in the intestinal cells and a decrease in the intensity of intestinal GFP expression compared with WT animals (Figure 6F, G). There was also a moderate decrease in the percentage of animals showing intestinal GFP expression in *smoc-1(tm7125)* mutants (Figure 6G). In contrast, nearly 100% of *lon-2(e678)* animals showed bright intestinal GFP expression, as compared to ∼92% for WT animals (Figure 6G). Collectively, these results suggest that *smoc-1* expression in the intestinal cells is positively regulated by BMP signaling.

### *smoc-1* functions cell non-autonomously to regulate body size and M lineage development

The *smoc-1* transcriptional reporters identified cells in the pharynx, intestine, and posterior hypodermis as *smoc-1*-expressing cells. To determine in which tissue(s) expression of *smoc-1* is sufficient to regulate BMP signaling, we used a set of promoters to drive *smoc-1* cDNA in a tissue-specific manner, and assayed for rescue of the *smoc-1(tm7125)* mutant phenotypes. Each rescuing construct was introduced into *smoc-1(tm7125)* worms for the body size assay, and into *smoc-1(tm7125); sma-9(cc604)* worms for the Susm assay.

As shown in Figure 7A, forced expression of *smoc-1* cDNA specifically within each individual *smoc-1*-expressing tissue (*ifb-2p* for intestinal cells (HUSKEN *et al.* 2008), *myo-2p* for pharyngeal muscles (OKKEMA *et al.* 1993), or *elt-3p* for hypodermal cells (GILLEARD *et al.* 1999)) not only rescued the small body size of *smoc-1(tm7125)* mutants, but also made the transgenic worms longer, just like *smoc-1* cDNA under the control of its own promoter. Forced expression of *smoc-1* cDNA in tissues that do not express *smoc-1* (*myo-3p* for BWMs (OKKEMA *et al.* 1993) or *rab-3p* for neurons (NONET *et al.* 1997)) also rescued the small body size of *smoc-1(tm7125)* mutants, and made the transgenic worms longer (Figure 7A). An exception is the lack of rescue of the body size phenotype in *smoc-1(tm7125)* mutants upon forced expression of *smoc-1* cDNA in the M lineage using the *hlh-8* promoter (HARFE *et al.* 1998). This could be due to the transient nature of *hlh-8* promoter activity in undifferentiated M lineage cells during larval development (HARFE *et al.* 1998).

**Fig 7.**
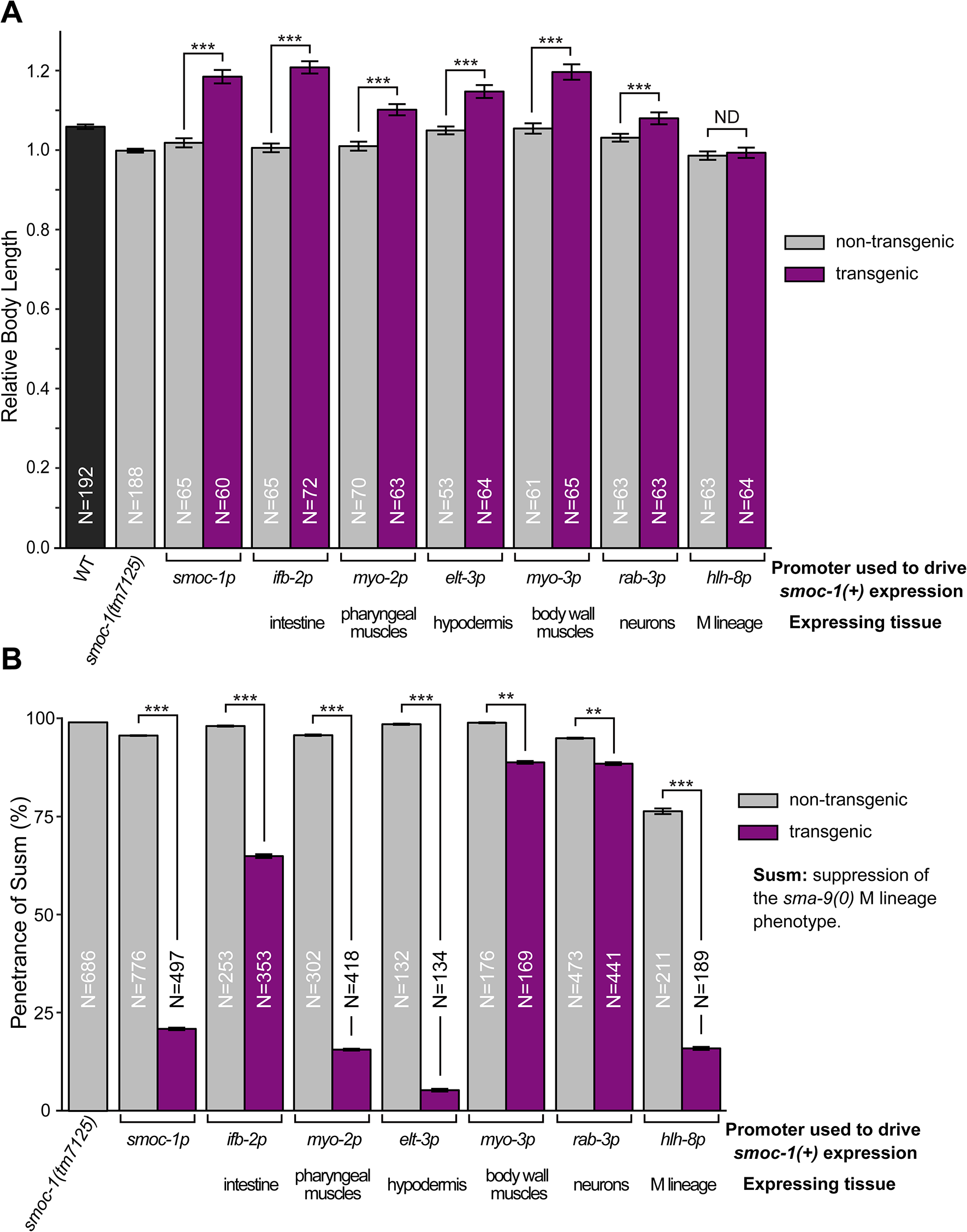
*smoc-1* functions cell non-autonomously to regulate body size and M lineage development. Tissue specific expression of *smoc-1* cDNA to test for rescue of the body size (**A**) or Susm (**B**) phenotype of *smoc-1(tm7125)* worms. *smoc-1* cDNA was driven by each specific promoter to allow expression in a given tissue. All constructs used the *unc-54* 3’UTR. For each construct, two independent transgenic lines were examined and the measurements were averaged. (**A**) Body sizes are relative to *smoc-1(tm7125)* mutant worms (set to 1.0), and all measurements were done on the same day. Error bars represent 95% CI. (**B**) The Susm phenotype was scored in the background of *smoc-1(tm7125); sma-9(cc604); CC::gfp*. Error bars represent standard error. ** *P*<0.001. *** *P*<0.0001. ND: no difference.

Similar to the body size rescue results, forced expression of *smoc-1* cDNA in both *smoc-1-* expressing cells (intestine, pharynx, or hypodermis) and cells that do not normally express *smoc-1* (BWMs, neurons, or the M lineage) rescued the Susm phenotype of *smoc-1(tm7125)* mutants (Figure 7B), although for reasons currently unknown, the rescuing efficiency appeared lower when *smoc-1* expression was forced in BWMs or neurons (Figure 7B). Taken together, our results demonstrate that SMOC-1 can function cell non-autonomously to regulate both body size and M lineage patterning. This is consistent with SMOC-1 being a putative secreted protein.

### Human SMOC proteins can partially rescue the *smoc-1(0)* mutant phenotype in *C. elegans*

As described above, SMOC-1 has two human homologs, SMOC1 (hSMOC1) and SMOC2 (hSMOC2). We next asked whether either of the human SMOCs can substitute for SMOC-1 function in *C. elegans*. We first generated plasmids by directly putting the coding region of hSMOC1 or hSMOC2 in between the 2kb *smoc-1* promoter and the *unc-54* 3’UTR (Table 2, Figure 8A), and tested their functionality using the Susm assay. Neither hSMOC1 nor hSMOC2 rescued the Susm phenotype of *smoc-1(tm7125)* worms (Figure 8B). We reasoned that the lack of rescue may be due to differences in the signal peptide between humans and *C. elegans*, causing the proteins to not be properly secreted from cells (TIAN *et al.* 2010). We next generated plasmids expressing chimeric SMOC proteins that have the worm SMOC-1 signal peptide (CelSP) followed by the extracellular region of hSMOC1 or hSMOC2 (Table 2, Figure 8A). Both CelSP::hSMOC1 and CelSP::hSMOC2 partially rescued the Susm phenotype of *smoc-1(tm7125)* mutants (Figure 8B), but failed to rescue the body size phenotype (Figure 8C). Nevertheless, these results demonstrate that CelSP::hSMOC1 and CelSP::hSMOC2 can function to regulate BMP signaling in a *C. elegans* trans-environment and suggest that the function of SMOC proteins in regulating BMP signaling is evolutionarily conserved from worms to humans.

**Fig 8.**
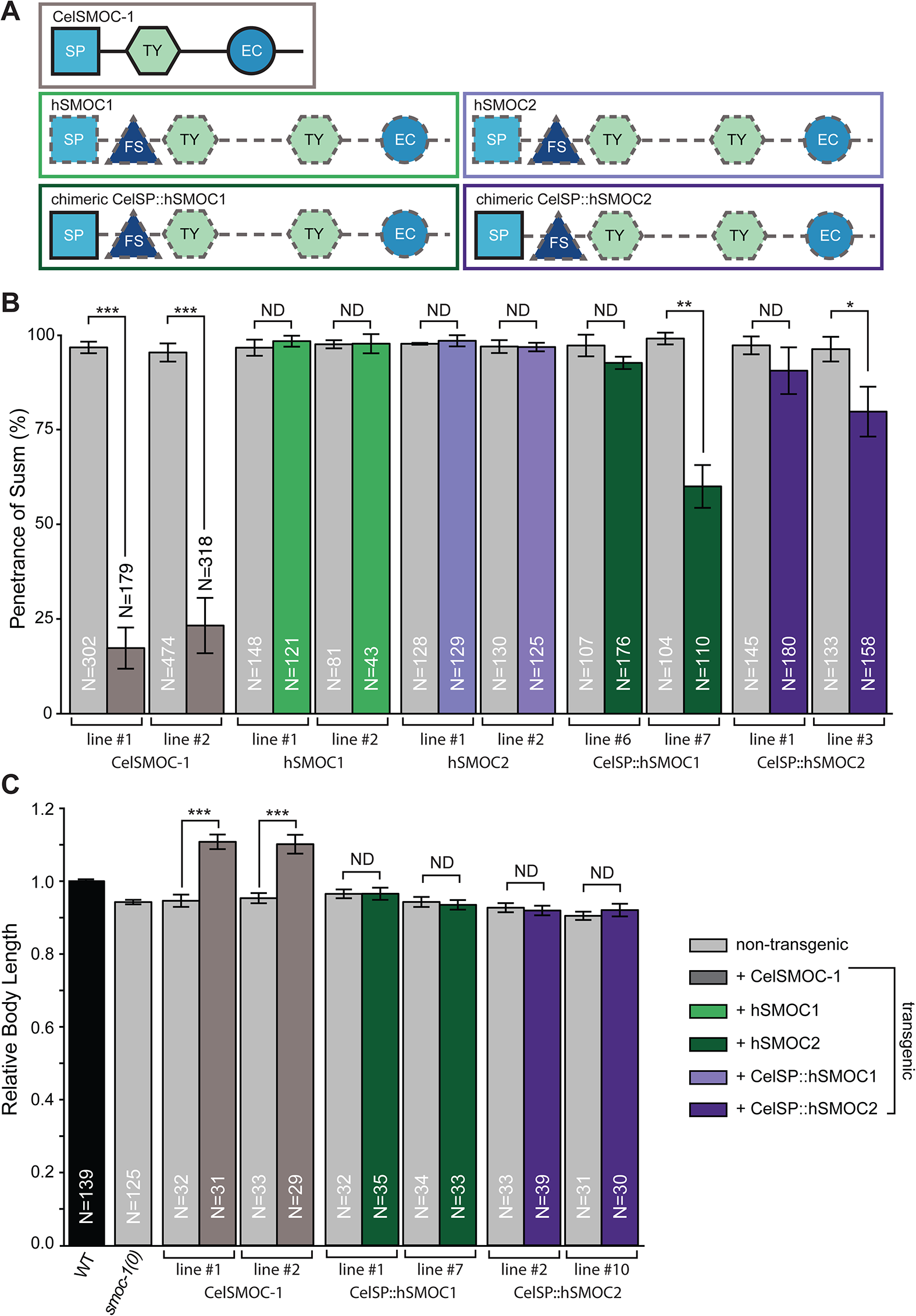
Human SMOC proteins can partially rescue the Susm phenotype of *smoc-1(0)* mutants. (**A**)Schematics of SMOC homologs tested for function in *C. elegans*. Solid black outline indicates *C. elegans* protein sequences. Dashed grey line indicates human protein sequences. All ORFs were cloned into the same vector with the same regulatory elements (2kb *smoc-1* promoter and *unc-54* 3’UTR), and each construct was tested for rescue of Susm (**B**) and body size (**C**) phenotype of *smoc-1(tm7125)* mutants. Two independent lines were assayed for each construct. The Susm phenotype was scored in the background of *smoc-1(tm7125); sma-9(cc604); CC::gfp*. Error bars represent standard error. (**C**) Body sizes are relative to WT worms (set to 1.0), and all measurements were done on the same day. Error bars represent 95% CI. * *P*<0.01. ** *P*<0.001. *** *P*<0.0001. ND: no difference.

## DISCUSSION

In this study, we identified SMOC-1, the sole *C. elegans* SMOC protein that belongs to the SPARC/BM40 family of matricellular proteins, as a key player in the BMP signaling pathway. *smoc-1(0)* mutants have a small body size and suppress the *sma-9(0)* M lineage defect, but *smoc-1(0)* mutants are not as small as null mutants in core components of the BMP pathway (Table 3, Figures 1, 3). These phenotypes resemble those caused by mutations in other modulators of the BMP pathway, such as DRAG-1/RGM (TIAN *et al.* 2010), TSP-21 (LIU *et al.* 2015), or SUP-17/ADAM10 (WANG *et al.* 2017), and are consistent with a modulatory role for SMOC-1 in the BMP pathway. Over-expression of *smoc-1* led to a significant increase in body size and an increase in RAD-SMAD reporter expression. Moreover, the long body size phenotype caused by *smoc-1(OE)* is completely suppressed by null mutations in the BMP ligand DBL-1 and the R-Smad SMA-3 (Figure 4). Collectively, these findings demonstrate that SMOC-1 functions through the BMP ligand DBL-1 and acts as a positive modulator to promote BMP signaling.

How might SMOC-1 function to promote BMP signaling? Our tissue specific rescue data coupled with the expression pattern of *smoc-1* (Figures 6, 7) showed that SMOC-1 functions cell non-autonomously to regulate BMP signaling. This is consistent with SMOC-1 being a predicted secreted protein. Strikingly, forced expression of *smoc-1* exclusively in pharyngeal muscles is sufficient to rescue both the body size and the Susm phenotype of *smoc-1(0)* mutants (Figure 7). Notably, the M lineage cells, where the Smad proteins function to regulate M lineage development (FOEHR *et al.* 2006), are located in the posterior of a developing larva, distant from the pharynx. Thus SMOC-1 can function over long distances, from a source located far from BMP-receiving cells, to regulate the output of BMP signaling.

The *Drosophila* homolog of SMOC-1, Pent, can also function over long distances to regulate Dpp/BMP signaling in the developing wing imaginal discs (VUILLEUMIER *et al.* 2010). In particular, Pent has been shown to bind to and induce the internalization of the BMP co-receptor Dally/glypican (a heparan sulfate proteoglycans (HSPG)), such that the trapping of Dpp/BMP by Dally is reduced, which in turn promotes the spreading of Dpp/BMP (NORMAN *et al.* 2016). Using a *Xenopus* animal cap transfer assay, Thomas and colleagues (THOMAS *et al.* 2017) showed that *Xenopus* SMOC-1 can also expand the range of BMP signaling by competing with BMP to bind to HSPGs. In *C. elegans*, the glypican homolog LON-2 is a known negative regulator of BMP signaling, and LON-2 can bind to BMP *in vitro* (GUMIENNY *et al.* 2007). LON-2/glypican has therefore been proposed to negatively regulate BMP signaling by sequestering the DBL-1/BMP ligand. Our genetic analysis between *lon-2(0)* and *smoc-1(0)* null mutations suggests that SMOC-1 antagonizes the function of LON-2 in regulating BMP signaling (Figure 5A). The phenotype of *smoc-1(0); lon-2(0)* double mutants is consistent with a model where SMOC-1 promotes BMP signaling by competing with DBL-1/BMP to bind LON-2/glypican. However, SMOC-1 must have LON-2/glypican-independent function(s), because *smoc-1(OE)* can further increase body size in the absence of LON-2/glypican, as in *smoc-1(OE); lon-2(0)* double mutants shown in Figure 5B.

The molecular mechanism underlying the LON-2/glypican-independent function of SMOC-1 is currently unknown. In addition to LON-2, there are five other HSPG-encoding genes in the *C. elegans* genome: *unc-52* (ROGALSKI *et al.* 1993; HALFTER *et al.* 1998; ACKLEY *et al.* 2001; RHINER *et al.* 2005; HRUS *et al.* 2007). It is possible that in addition to LON-2/glypican, one or multiple of these other HSPGs also functions with SMOC-1 to regulate BMP signaling. Alternatively, SMOC-1 may promote BMP signaling by interacting with other cell surface or extracellular BMP regulators or even with DBL-1/BMP itself to promote BMP signaling. Any LON-2/glypican-independent function of SMOC-1 still requires DBL-1/BMP, because *smoc-1(OE); dbl-1(0)* double mutants are as small as *dbl-1(0)* null mutants. Our model proposing SMOC-1 has dual modes of action to regulate BMP signaling is consistent with structure-function analysis of *Xenopus* SMOC-1 (XSMOC-1), whose EC domains can bind to HSPG and promote BMP spreading, while the TY domains are necessary for XSMOC-1 to inhibit BMP signaling (THOMAS *et al.* 2017). We have shown that both the TY domain and the EC domain in *C. elegans* SMOC-1 are important for its function in BMP signaling, because mutations in either domain disrupt the function of SMOC-1 (Figure 2). Further dissection of the roles of each of these domains at the molecular level will help clarify the mechanisms underlying SMOC-1 function in the BMP pathway.

In this study, we have shown that in addition to being a positive regulator of BMP signaling, *smoc-1* is also positively regulated by BMP signaling at the transcriptional level (Figure 6). Whether *smoc-1* is directly or indirectly regulated by BMP signaling remains to be determined. Nevertheless, our results suggest a model in which SMOC-1 functions in a positive feedback loop to regulate BMP signaling (Figure 9). Whether or not *smoc-1* expression is directly regulated by BMP signaling is currently unknown.

**Fig 9.**
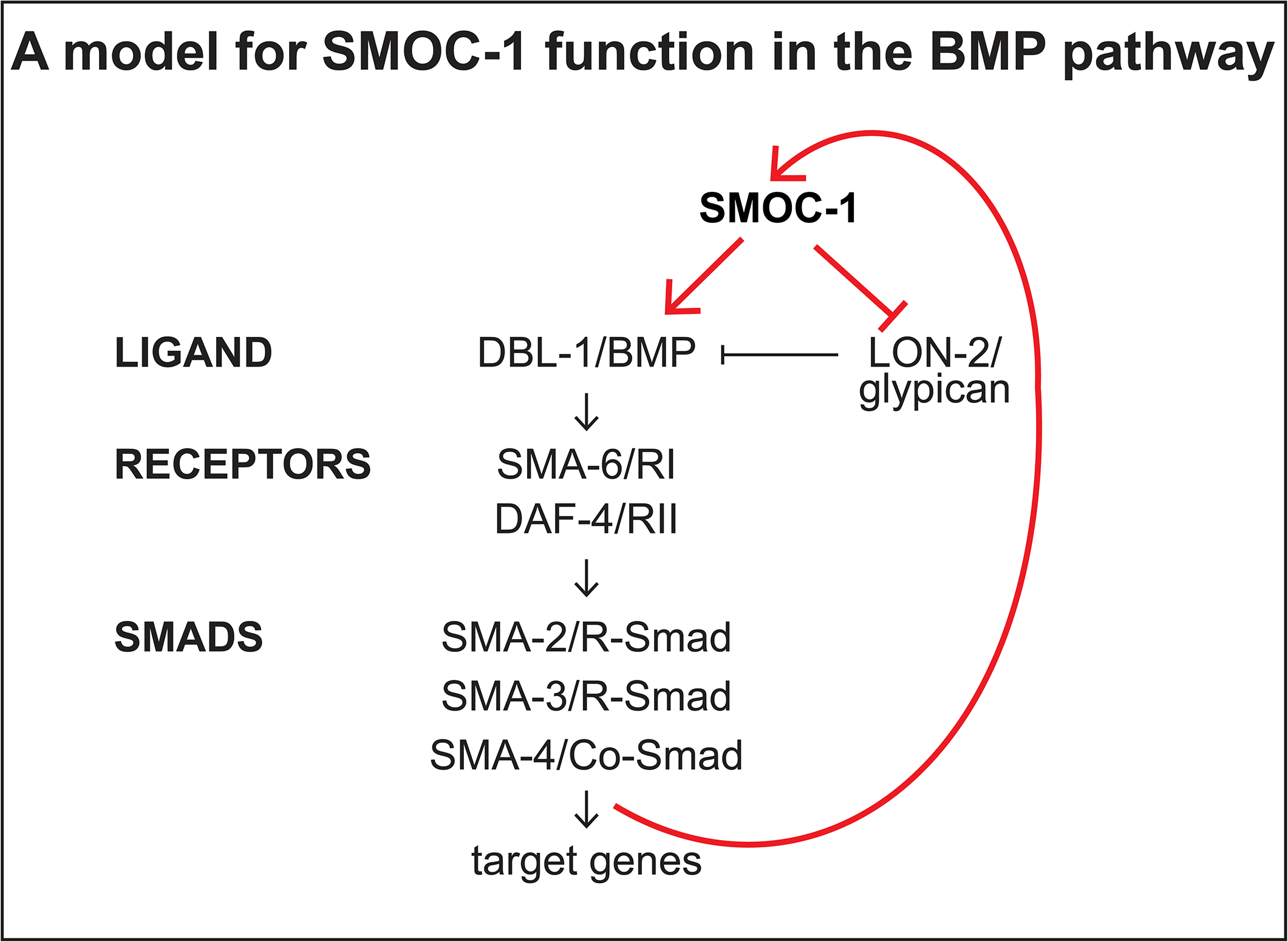
A model for SMOC-1 function in the BMP pathway. SMOC-1 acts through the BMP ligand DBL-1/BMP, and in part by antagonizing LON-2/glypican, to promote BMP signaling. BMP signaling in turn promotes the intestinal expression of *smoc-1,* thus creating a positive feedback loop.

In addition to their roles in regulating BMP signaling, SMOC proteins can also function in other signaling pathways. Pent has been shown to play a role in regulating Wg signaling in the *Drosophila* wing (NORMAN *et al.* 2016). Human SMOC1 can bind to the TGFβ co-receptor endoglin to regulate TGFβ signaling in endothelial cells (AWWAD *et al.* 2015), while SMOC2 can potentiate endothelial growth factor or fibroblast growth factor activity to promote angiogenesis in cultured human umbilical vein endothelial cells (HUVECs) (ROCNIK *et al.* 2006). Our genetic analysis suggests that SMOC-1 does not play a key role in regulating the TGFβ-like dauer pathway (Table 4). Whether SMOC-1 is involved in other signaling pathways in *C. elegans* is currently unknown.

There are two SMOC homologs in mammals. SMOC1 is essential for eye and limb development in mice, and mutations in SMOC1 in humans cause microphthalmia with limb anomalies (MLA) and ophthalmo-acromelic syndrome (OAS) (also known as Waardenburg anophthalmia syndrome (WAS)), both of which affect eye and limb development (OKADA *et al.* 2011; RAINGER *et al.* 2011). Mutations in hSMOC2 have also been found to be associated with defects in dental development (BLOCH-ZUPAN *et al.* 2011; ALFAWAZ *et al.* 2013) and vitiligo (ALKHATEEB *et al.* 2010; BIRLEA *et al.* 2010). QTL mapping in different dog breeds have found that a retrotransposon insertion that disrupts SMOC2 splicing and reduces its expression is associated with canine brachycephaly (MARCHANT *et al.* 2017). In addition, several different types of brain tumors exhibit altered expression of SMOC1 (BRELLIER *et al.* 2011), while SMOC2 is an intestinal stem cell signature gene (MUNOZ *et al.* 2012) that is required for L1-mediated colon cancer progression (SHVAB *et al.* 2016). Notably, BMP signaling is known to play important roles in eye, tooth and limb development, and abnormal BMP signaling can cause cancer (THAWANI *et al.* 2010). Here, we have demonstrated that both hSMOC1 and hSMOC2 can partially rescue the Susm phenotype of *smoc-1(0)* mutants (Figure 8), suggesting that the function of SMOC proteins in regulating BMP signaling is evolutionarily conserved. Future studies on how SMOC-1 functions to regulate BMP signaling in an *in vivo* system such as *C. elegans* may have implications for human health.

## ACKNOWLEDGMENTS

We thank Oliver Hobert and Shohei Mitani for plasmids or strains, Florencia Schlamp for advice on statistical analysis using R, Gabriela Rojas for help generating the *smoc-1(OE); RAD-SMAD* strains, and the rest of the Liu lab for critical comments and suggestions. Some strains were obtained from the *C. elegans* Genetics Center, which is funded by NIH Office of 27 Research Infrastructure Programs (P40 OD010440). This study was supported by the National Institutes of Health grants R01 GM066953 and R01 GM103869 to JL. MSD was partially supported by a NIH Predoctoral Training Grant (T32GM007617), a Dean’s Excellence Fellowship by the Cornell Graduate School, and an NSF Graduate Research Fellowship (DGE-1650441). AE was a student in the MBG-REU program, which was supported by the NSF (DBI1659534), Department of Molecular Biology and Genetics, Weill Institute of Cell and Molecular Biology, and Division of Nutritional Sciences at Cornell University. ANM was a Hunter R. Rawlings III Presidential Research Scholar at Cornell University.

